# Flagellar targeting of an arginine kinase requires a conserved lipidated intraflagellar transport (LIFT) pathway in *Trypanosoma brucei*

**DOI:** 10.1101/2020.05.07.083568

**Authors:** Maneesha Pandey, Yameng Huang, Teck Kwang Lim, Qingsong Lin, Cynthia Y. He

## Abstract

Both intraflagellar transport (IFT) and lipidated intraflagellar transport (LIFT) pathways are essential for cilia/flagella biogenesis, motility and sensory functions. In the LIFT pathway, lipidated cargoes are transported into the cilia through the coordinated actions of cargo carrier proteins such as Unc119 or PDE6δ, as well as small GTPases Arl13b and Arl3 in the cilium. Our previous studies revealed a single Arl13b ortholog in the evolutionarily divergent *Trypanosoma brucei*. TbArl13 catalyses two TbArl3 homologs, TbArl3A and TbArl3C, suggesting the presence of a conserved LIFT pathway in these protozoan parasites. Only a single homolog to the cargo carrier protein Unc119 was identified in *T. brucei* genome, but its function in lipidated protein transport has not been characterized. In this study, we exploited the proximity-based biotinylation approach to identify binding partners of TbUnc119. We showed that TbUnc119 binds to a flagellar arginine kinase TbAK3 in a myristoylation-dependent manner and is responsible for its targeting and enrichment in the flagellum. Interestingly, only TbArl3A, but not TbArl3C interacts with TbUnc119 in a GTP-dependant manner, suggesting functional specialization of Arl3-GTPases in *T. brucei*. This study establishes the function of TbUnc119 as a myristoylated cargo carrier and supports the presence of a conserved LIFT pathway in *T. brucei.*

## Introduction

Cilia or eukaryotic flagella are present in eukaryotic organisms ranging from protists, invertebrates to vertebrates. Depending on their structure and protein compositions, cilia and flagella can perform sensory functions or impart motility. Ciliary mutations and malfunctioning have been implicated in many diseases collectively known as ciliopathies (1).

Ciliary proteins are synthesized in the cytosol and trafficked to the ciliary compartment by two main pathways, the intraflagellar transport (IFT) and the lipidated intraflagellar transport (LIFT) (1,2). IFT has been extensively characterized with well-documented functions in anterograde and retrograde transport of ciliary structural components. LIFT, on the other hand, has recently emerged as a parallel trafficking pathway dedicated to lipidated cargoes associated with ciliary membrane (3-6). Only a few lipidated proteins have been identified as LIFT cargoes, and most of these proteins are important for ciliary signalling functions (5-7).

In mammals, LIFT comprises of small GTPases Arl13b and Arl3, Arl3-GTPase activating protein RP2, as well as carrier proteins Phosphodiesterase 6δ (PDE6δ) and Unc119 paralogs (2). PDE6δ or Unc119 binds to lipidated cargoes synthesized in the cytosol and facilitates their import into the ciliary lumen. Arl13b is enriched in the cilia, where it acts as a guanine nucleotide exchange factor (GEF) on Arl3 (3). Activated, GTP-bound Arl3 can bind to PDE6δ or Unc119, and functions as a displacement factor to release lipidated cargoes associated with the carrier proteins inside of the ciliary lumen (8-10). Arl3-GTP hydrolysis is then catalysed by RP2 at the base of the cilium, where Arl3-GDP dissociates from the carrier proteins (11).

PDE6δ and Unc119 paralogs contain a conserved phosphodiesterase domain, which is crucial for interaction with lipidated cargoes (12). PDE6δ binds ciliary farnesylated proteins such as inositol polyphosphate 5’-phosphatase E (INPP5E) (7) and is required for its ciliary targeting. PDE6δ also interacts with non-ciliary cargoes such as prenylated RAS-GTPases and affects their membrane distribution and signalling functions (13). Unc119 paralogs Unc119A and Unc119B, on the other hand, share 60% sequence homology and both carry myristoylated proteins into the cilia (12). Only a few myristoylated cargoes have been identified (5), and Nephrocystin-3 (NPHP3) is the only ciliary cargo identified till date that interacts with both Unc119A and Unc119B (6). Like PDE6δ, Unc119A has also been shown to have non-ciliary functions via interactions with Src-type tyrosine kinases Lyn (14), Fyn (15), and Lck as well as Rab11 (16,17) and Dynamin GTPases (18), all of which have predicted N-myristoylation sites. Through these interactions, Unc119A influences the distribution and signalling functions of these proteins.

The function of Unc119 has also been extensively characterized in *C. elegans*, where it is also a lipid binding protein required for G protein trafficking in sensory neurons (19). Additionally, a recent study has shown that *C. elegans* Unc119 interacts with both Arl3 and Arl13, stabilizing the interaction between Arl3 and Arl13, and facilitating GTP activation of Arl3 (20). Importantly, *C. elegans* Unc119 binds to Arl3 independent of its GTP-bound state, making Unc119 an unlikely effector of Arl3-GTP. The *C. elegans* Unc119 thus functions differently to its mammalian counterparts, which may represent functional divergence in lower ciliated organisms (20).

*Trypanosoma brucei*, causative agent of human African trypanosomiasis (sleeping sickness) as well as nagana in domestic animals, is a protozoan parasite belonging to the Kinetoplastid group, which are considered as one of the earliest-divergent eukaryotic organisms (21). *T. brucei* is also emerging as a useful model to understand flagellar structure, biogenesis and functions (22). The flagellum of *T. brucei* has both signalling and motility functions (23) and is crucial for the viability and pathogenesis of this parasite. Both IFT and LIFT pathway components have been identified in *T. brucei* (22,24). While the function and regulation of the IFT pathway has been extensively characterized (22), the presence of a conserved LIFT pathway in *T. brucei* was only recently recognized. A single orthologue of Arl13b was found in *T. brucei* genome, with its protein product enriched in the flagellar axoneme via a Docking and Dimerization domain (24). Interestingly, *T. brucei* has three Arl3 homologs namely TbArl3A (Tb927.3.3450), TbArl3B (Tb927.10.8580) and TbArl3C (Tb927.6.3650). TbArl13 interacts and catalyses nucleotide exchange on both TbArl3A and TbArl3C, but not TbArl3B. Consistently, only TbArl3A and TbArl3C exhibit flagellar biogenesis effects upon overexpression of the GTP-locked mutants (24).

A single Unc119 ortholog (TbUnc119, Tb927.2.4580) was identified in an in silico screen of *T. brucei* genome (25). TbUnc119 is present in the flagellum, but depletion of TbUnc119 via tetracycline-inducible RNA interference (RNAi) does not produce any observable effect on cell growth or motility (25). Thus, the cellular function of TbUnc119 is not known and its role in lipidated protein transport in *T. brucei* has not been studied. In this study we used BioID, a proximity-based biotinylation method (26,27) to identify interacting partners of TbUnc119. Our results identified a flagellar arginine kinase TbAK3 as a TbUnc119 cargo. We also showed that TbArl3A but not TbArl3C binds to Unc119, emphasizing the functional difference between different TbArl3-GTPase isoforms in *T. brucei*.

## Results

### Kinetoplastids contain a single Unc119 homolog

*Trypanosoma brucei* has a single Unc119 homolog (TbUnc119) encoded by Tb927.2.4580, which has been shown to be a flagellar protein in an earlier study (25). Phylogenetic analyses were then performed on TbUnc119 and Unc119/PDE6δ homologs identified in various model organisms (Fig. S1A). TbUnc119 formed a clad with other Unc119 homologs distinct from PDE6δ proteins. Unc119 is highly conserved among kinetoplastids (Fig. S1B). Notably, PDE6δ homologue could not be found despite extensive searches of the *T. brucei* genome. Further searches using NCBI BLAST confirmed the absence of PDE6δ in all Kinetoplastid members and most single cellular eukaryotes we have examined, with the possible exception of *Paramecium tetraurelia* (28). Together these results suggest that Unc119 is likely the only conserved lipidated protein carrier belonging to the Unc119 supergene family (28) in *T. brucei* and other Kinetoplastid organisms.

The knockdown of TbUnc119 did not produce detectable growth defects in the insect-infectious procyclic form (PCF) cells (Fig. S2A), which corroborates the previous study (25). In the mammal infectious bloodstream form (BSF) *T. brucei*, a mild growth delay was consistently observed post TbUnc119-RNAi induction (Fig. S2B). Similar to TbUnc119-silencing in the PCF cells, no significant phenotypic changes were observed in the BSF cells. Taken together, these results confirmed that TbUnc119 is not essential for procyclic and BSF cell survival or flagellar biogenesis in culture. These results are also consistent with the non-lethal mutant phenotypes of Unc119 orthologues previously reported in *C. elegans* (29) or zebrafish (30).

### Identification of TbUnc119-interacting proteins by proximity-based biotinylation

In both *C. elegans* and mammals, Unc119 is characterized as a cargo carrier/chaperone involved in intraflagellar transport of myristoylated ciliary proteins, and Unc119 association with the cargo is regulated by the Arl13b-Arl3 pathway (3,6). *C. elegans* Unc119 additionally functions in stabilizing the Arl13b-Arl3 interaction, suggesting functional divergence of Unc119 in lower eukaryotes (20). The function of the LIFT pathway and its flagellar cargoes have never been examined in the evolutionarily-divergent *T. brucei*, we therefore decided to revisit the function of TbUnc119, by investigating its interacting proteins.

We utilized the proximity-dependant biotinylation approach, using an improved version of biotin ligase BioID2 (26,31). The BioID2 tag was fused to either the N-terminus (3HA-BioID2-TbUnc119) or the C-terminus (TbUnc119-BioID2-HA) of TbUnc119 and expressed using a cumate-inducible expression system in the procyclic cells (32). Both 3HA-BioID2-TbUnc119 and TbUnc119-BioID2-HA cells showed strong labelling throughout the cytoplasm (Fig. S3), by anti-HA that stains the TbUnc119 fusions and streptavidin-Alexa Fluor 568 that stains the biotinylated products. Weak signal was also observed along the flagellum (Fig. S3B and S3C, arrowheads). These results suggest both flagellar and cytoplasmic presence of TbUnc119 and its biotinylated products. Similar cytoplasmic presence is also apparent in cells with endogenous expression of mNeonGreen-tagged TbUnc119, either at the N- or C-terminus (http://tryptag.org/?query=Tb927.2.4580) (33). Biotinylated proteins from both 3HA-BioID2-TbUnc119 and TbUnc119-BioID2-HA cells were affinity purified (Fig. S3, D and E) and analysed by LC/MS-MS. A total of 138 and 43 candidates from 3HA-BioID2-TbUnc119 and TbUnc119-BioID2-HA cells, respectively, were identified (Fig. 1A). Among them, 32 high-confidence candidates were found in both 3HA-BioID2-TbUnc119 and TbUnc119-BioID2-HA cells (Fig. 1B; Table S1). It is not clear why 3HA-BioID2-TbUnc119 had more hits identified than TbUnc119-BioID2-HA cells. One possibility is that the position of the BioID2 tag at the C-terminus of TbUnc119 may interfere with its interaction with other proteins.

**Fig. 1.**
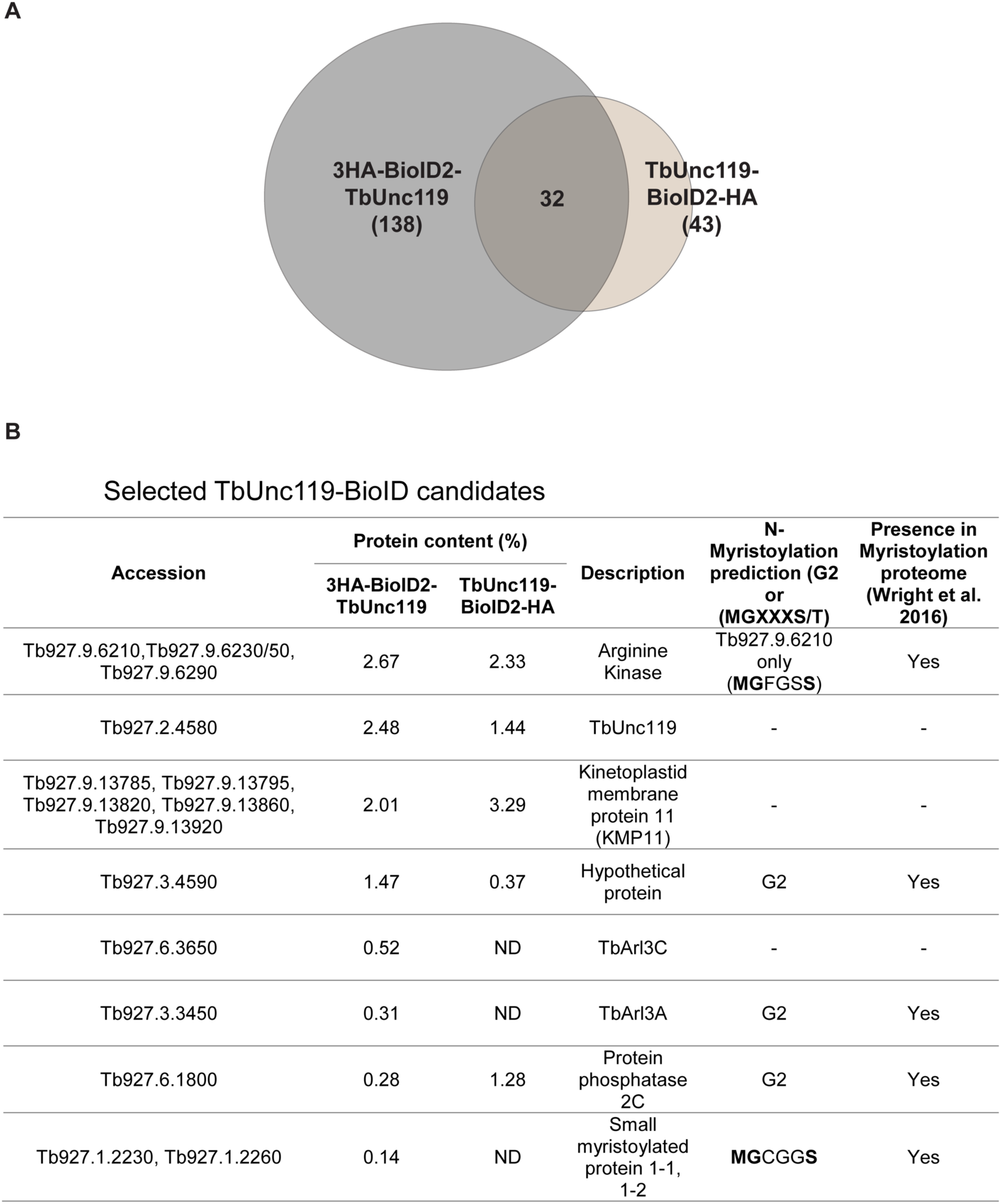
Proximity-dependent biotinylation screening for TbUnc119 interacting proteins. (**A**) A Venn diagram summarizing the mass spectrometry results of two independent BioID experiments, one using cells stably expressing 3HA-BioID2-TbUnc119 with BioID2 fused to the N-terminus of TbUnc119 and the other TbUnc119-BioID2-HA with BioID2 fused to the C-terminus. (**B**) List of selected BioID candidates. Protein candidates identified are ordered according to calculated protein content in the BioID experiment using 3HA-BioID2-TbUnc119 as the bait. Protein content of the candidates in the TbUnc119-BioID2-HA experiment is also shown wherever applicable. The description of the protein was obtained from the Kinetoplastid Genomics resource (www.tritrypdb.org). ND: Not Detected.

A group of arginine kinases (AKs) and kinetoplastid membrane proteins, KMP-11, were found to top the list (Fig. 1B). Of the three highly-similar AK proteins identified, only TbAK3 (encoded by Tb927.9.6210 (34); and named AK1 in another study (35) contains a myristoylation consensus sequence. TbAK3 was also identified as a high-confidence candidate in a recent myristoylation proteomics study (36). Small myristoylated proteins (TbSMP1-1 encoded by Tb927.1.2230 and TbSMP1-2 by Tb927.1.2260), which also possess a consensus myristoylation sequence were identified in the 3HA-BioID2-TbUnc119 list. These kinetoplastid-specific proteins resemble calpain like proteins and are associated with the cell membrane (37). TbArl3A and TbArl3C, both components of *T. brucei* flagellum with confirmed flagellar functions (24), were also identified in 3HA-BioID2-TbUnc119 BioID.

### The flagellar targeting of TbAK3 requires TbUnc119

Three Arginine Kinases are found in *T. brucei*, sharing 85-99% sequence identity (35) and thus could not be distinguished in the BioID mass spectrometry results. TbAK1 (encoded by Tb927.9.6290) is localized throughout the cytoplasm; and TbAK2 (encoded by Tb927.9.6250) is associated with the glycosomes, a peroxisome-like organelle in *T. brucei (35)*. TbAK3 (encoded by Tb927.9.6210) is present on the flagellar membrane and is the only TbAK that is myristoylated (35,36,38).

To test if TbAK3 interacted with TbUnc119, TbAK3 was fused to a small BB2 tag (39) at the C-terminus and co-expressed with GFP-TbUnc119. Immunoprecipitation was then performed using GFP nAb-conjugated beads. TbAK3-BB2 co-immunoprecipitated with GFP-TbUnc119, but not GFP only (Fig. 2A). As another control, TbAK1-BB2 did not co-immunoprecipitate with GFP-TbUnc119, suggesting specific interaction between TbAK3 and TbUnc119. The TbAK3-TbUnc119 interaction is myristoylation-dependent, as TbAK3(G2A)-BB2 failed to co-immunoprecipitate with GFP-TbUnc119. The myristoylation mutation also disrupted the flagellar localization of TbAK3 (Fig. 2, B and C). Together these results indicate that the flagellar localization of TbAK3 and its interaction with TbUnc119 are both myristoylation-dependent.

**Fig. 2.**
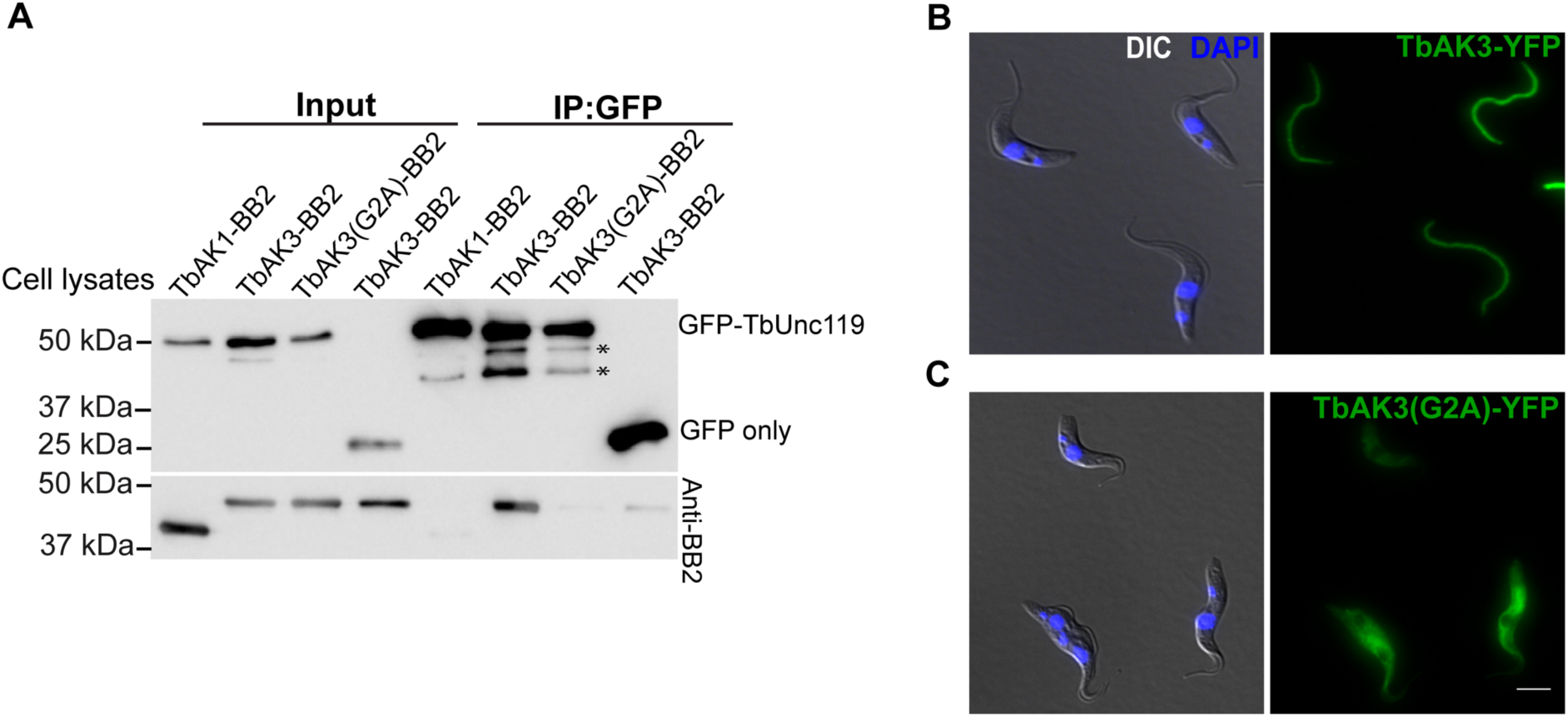
Myristoylation of TbAK3 is critical for its flagellar targeting and its interaction with TbUnc119. (**A**) Lysates of cells stably expressing GFP-TbUnc119 and TbAK1-BB2, TbAK3-BB2 or TbAK3(G2A)-BB2, respectively, were incubated with GFP-nAb beads. Proteins bound to the beads were fractionated on SDS-PAGE followed by immune-blotting with anti-GFP and anti-BB2 antibodies. Cells co-expressing GFP and TbAK3-BB2 were used as a negative control. Asterisks indicate possible degradation products of GFP-TbUnc119. Input: 3% of cell lysates. (**B, C**) Cells expressing TbAK3-YFP or TbAK3(G2A)-YFP were viewed after fixation with 4% PFA. Nuclear and kinetoplast DNA were stained with DAPI (blue). Scale bar: 5 μm.

Next we asked whether TbAK3 flagellar targeting required TbUnc119. We tagged one endogenous allele of TbAK3 with fluorescent reporter mNeonGreen in the TbUnc119-RNAi cell line. In control cells without TbUnc119-RNAi induction, TbAK3-mNeonGreen was enriched in the flagella with weak signal in the cytosol (Fig. 3A). Upon TbUnc119-RNAi induction, the flagellum enrichment of TbAK3 diminished and increased cytosolic signal was observed (Fig. 3A, quantitated in Fig. 3B). To further establish the flagellar targeting dynamics of TbAK3, we generated a stable cell line with tetracycline-inducible TbUnc119-RNAi and cumate-inducible expression of TbAK3-BB2. The expression of TbAK3-BB2 was induced for a fixed period of 24 hours, at different times post TbUnc119-RNAi induction. A gradual reduction in flagellar TbAK3 and an increase in the cytoplasmic TbAK3 was observed over the course of TbUnc119-RNAi (Fig. 3C, quantitated in 3D), despite similar expression levels of TbAK3 at different timepoints (Fig. 3E). These results suggested that the loss of TbUnc119 inhibited the entry of TbAK3 into the flagellum.

**Fig. 3.**
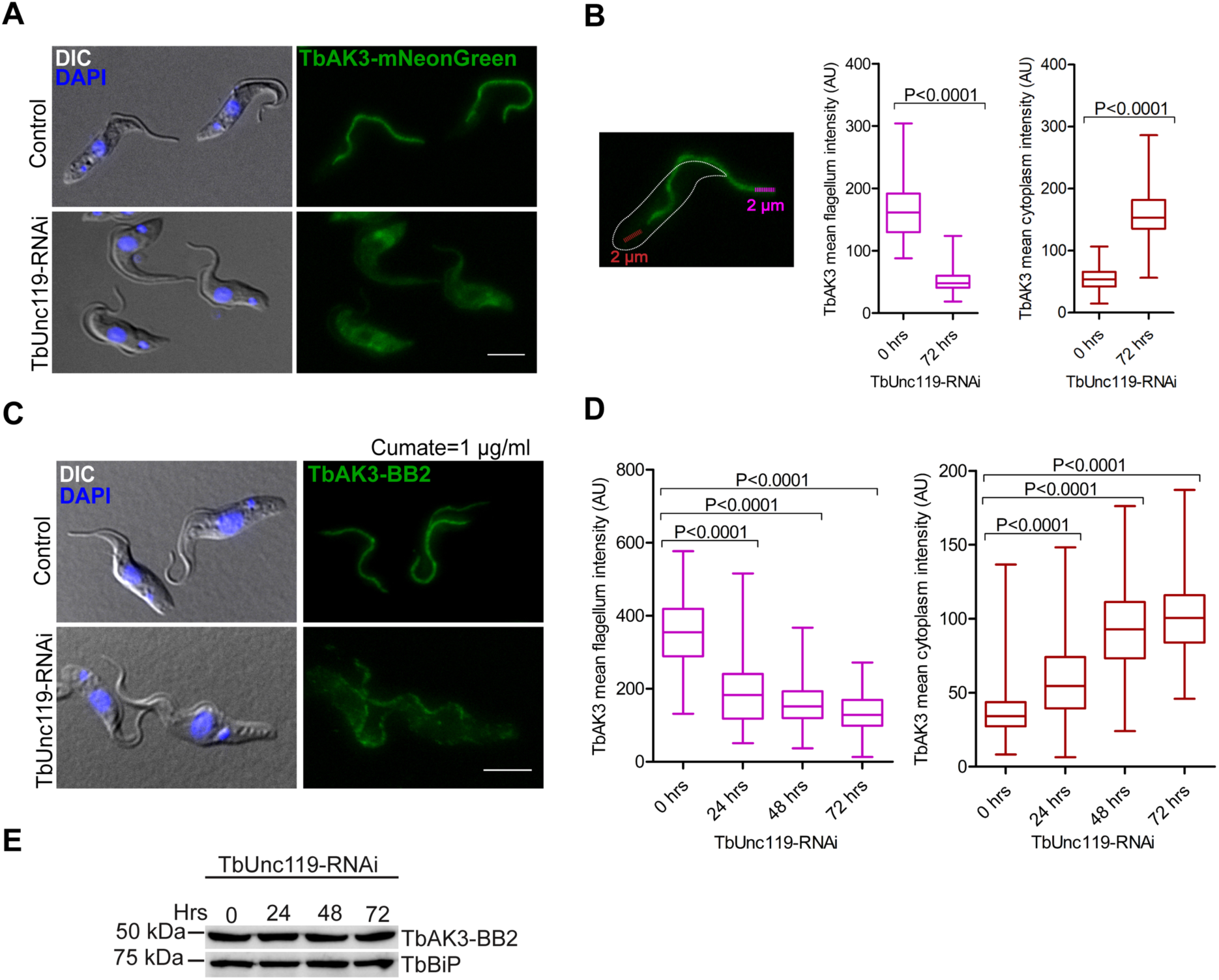
Flagellar targeting of TbAK3 requires TbUnc119. (**A**) Procyclic cells stably expressing TbAK3-mNeonGreen from an endogenous allele were induced for TbUnc119-RNAi. Control and induced cells were fixed with 4% PFA. (**B**) TbAK3-mNeonGreen intensity in the flagellum and in the cytosol was measured using plot profile function as illustrated here and detailed in the experimental procedures. Results are shown as box plots, with the whiskers marking the minimum and maximum values, the box showing the 25th to 75th percentiles and the bars in the box showing the median. N>150 cells were measured for each condition at each time point. Two-tailed student’s t-test was performed, and p values are indicated in the plots. (**C, D, E)** Cells containing tetracycline-inducible TbUnc119-RNAi and cumate-inducible TbAK3-BB2 expression were induced for TbUnc119-RNAi for 0, 24 or 48 hours prior to induction of TbAK3-BB2 expression. The induction of TbAK3-BB2 expression was fixed at 1µg/ml cumate for 24 hours to ensure similar expression levels in different experiments (E). Cells were then fixed and processed for immunostaining with anti-BB2 (C). Quantitation of TbAK3-BB2 signal in the flagellum and the cytosol was performed as illustrated in (B). N>100 cells were measured for each condition at each time point (D). Scale bar: 5 µm.

### TbUnc119 binds to TbSMP1-1, but is not required for TbSMP1-1 intracellular distribution

To address whether TbUnc119 may have non-ciliary functions as observed with animal Unc119 orthologs, we examined TbUnc119-BioID candidates for non-ciliary myristoylated proteins. The small myristoylated protein TbSMP1-1 (encoded by Tb927.1.2230) contains an N-terminal myristoylation site (G2) and is enriched at the cell membrane of *T. brucei* (37). The interaction between TbSMP1-1 and TbUnc119 was confirmed by co-immunoprecipitation (Fig. 4A). While TbSMP1-1-GFP was enriched in cell periphery consistent with plasma membrane association, TbSMP1-1(G2A)-GFP mutant lost cell membrane enrichment and was found throughout the cytosol (Fig. 4, B and C). This observation was further confirmed by profiling the fluorescent intensity across randomly selected TbSMP1-1-GFP and TbSMP1-1(G2A)-GFP expressing cells (Fig. 4, B and C). Silencing of TbUnc119 however, had no detectable effects on TbSMP1-1-GFP distribution in the cell (Fig. 4D).

**Fig. 4.**
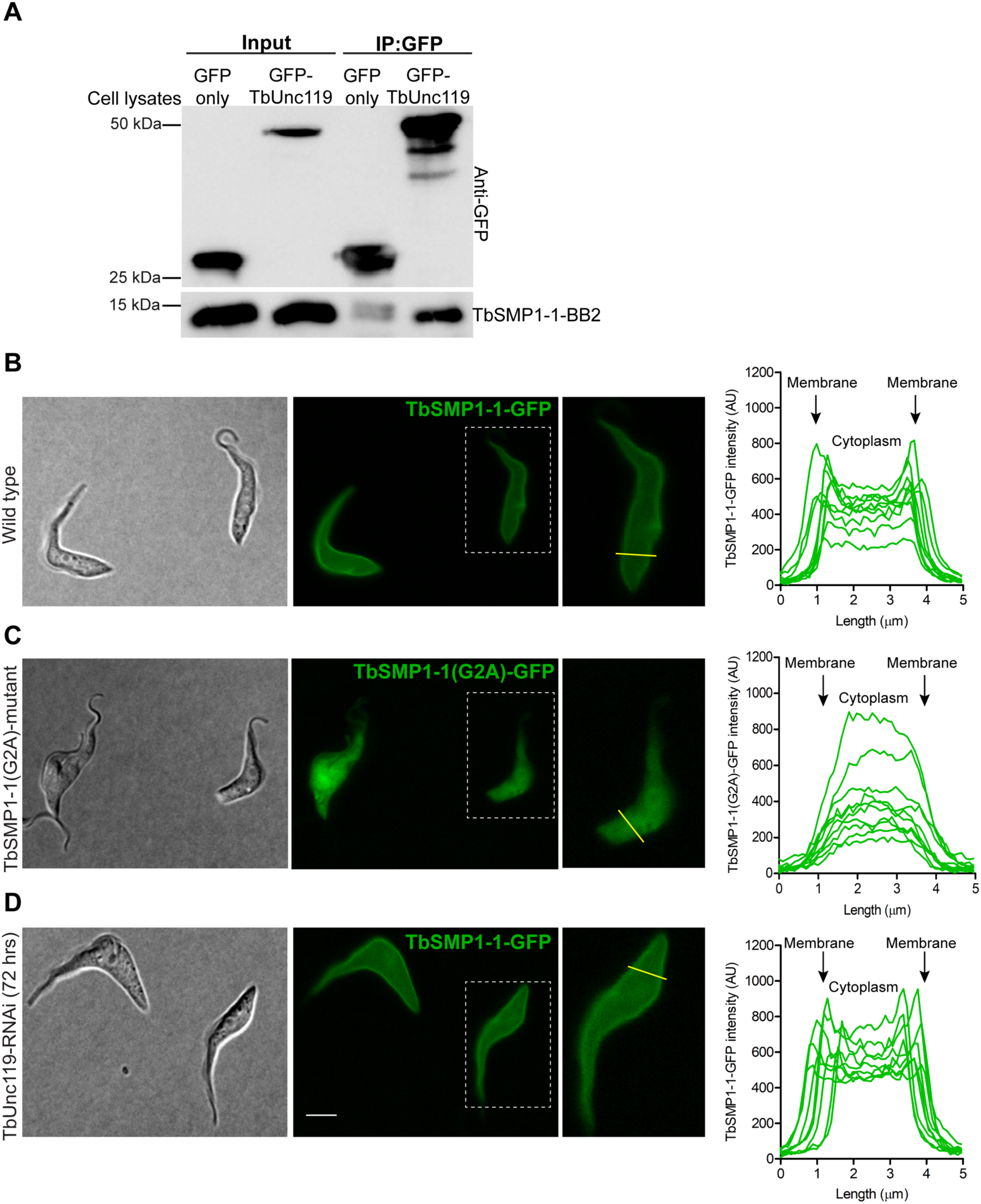
The cell membrane association of TbSMP1-1 is myristoylation-dependent but TbUnc119-independent. (**A**) TbSMP1-1-BB2 co-immunoprecipitated with GFP-TbUnc119, but not GFP only. (**B, C, D**) Cells expressing TbSMP1-1-GFP (B) or TbSMP1-1(G2A)-GFP (C) were immobilized on agarose gel and imaged live, to best visualize the plasma membrane association of TbSMP1-1-GFP. The intracellular distribution of TbSMP1-1-GFP was also monitored in live cells induced for TbUnc119-RNAi (D). All GFP images were collected at constant exposure time. The distribution of TbSMP1-1-GFP in control and TbUnc119-RNAi cells was measured using plot profiling. A line of 5µm length was drawn across the entire cell body, encompassing cell membranes at both ends, and fluorescence intensity along the length of this line was plotted. 10 representative cells were shown for control and TbUnc119-RNAi cells. Insets show enlarged images of a representative cell from each sample that were selected for intensity measurements. Scale bar: 5 µm.

### TbUnc119 interacts specifically with Arl3A in a GTP-dependent manner

The best characterized function of Unc119 is in the context of the LIFT pathway as a carrier for myristoylated cargoes. Once the cargo-Unc119 complex is inside of the ciliary lumen, Arl3-GTP acts as a displacement factor, binds to Unc119 and releases the cargo (4,6). Unlike vertebrates that contain only a single Arl3 protein, *T. brucei* has two Arl3 homologues, TbArl3A and TbArl3C, that are both associated with the flagellum and exhibit flagellar phenotypes when overexpressed as GTP-locked forms (24). Interestingly, both TbArl3A and TbArl3C were found in the TbUnc119 BioID screen, albeit only in the 3HA-BioID2-TbUnc119 cells.

To examine the interaction specificity of TbUnc119 with TbArl3-GTPases, co-immunoprecipitation assays were performed on cells stably expressing GFP-TbUnc119 and mNeonGreen-tagged TbArl3A or TbArl3C from an endogenous allele. TbArl3A-mNeonGreen-BB2 specifically co-precipitated with GFP-TbUnc119, but not with GFP only (Fig. 5A); TbArl3C-mNeonGreen-BB2 did not co-precipitate with either GFP-TbUnc119 or GFP alone (Fig. 5B). Direct interaction between TbArl3A and TbUnc119 was confirmed by pulldown assays using purified His-TbUnc119 and GST-TbArl3A (Fig. 5, C and D). Importantly, TbUnc119 did not interact with TbArl13 in the co-immunoprecipitation assays (Fig. 4A, also see Fig. 6C below). This is distinct to the Arl13-Arl3-Unc119 mutual interactions observed in *C. elegans* (20). Additionally, TbUnc119 did not interact with TbArl2 (encoded by Tb927.10.4250) (Fig. S4), which exhibits high sequence similarities to TbArl3A and TbArl3C (24,40). These interaction results are summarized in Fig. 5E.

**Fig. 5.**
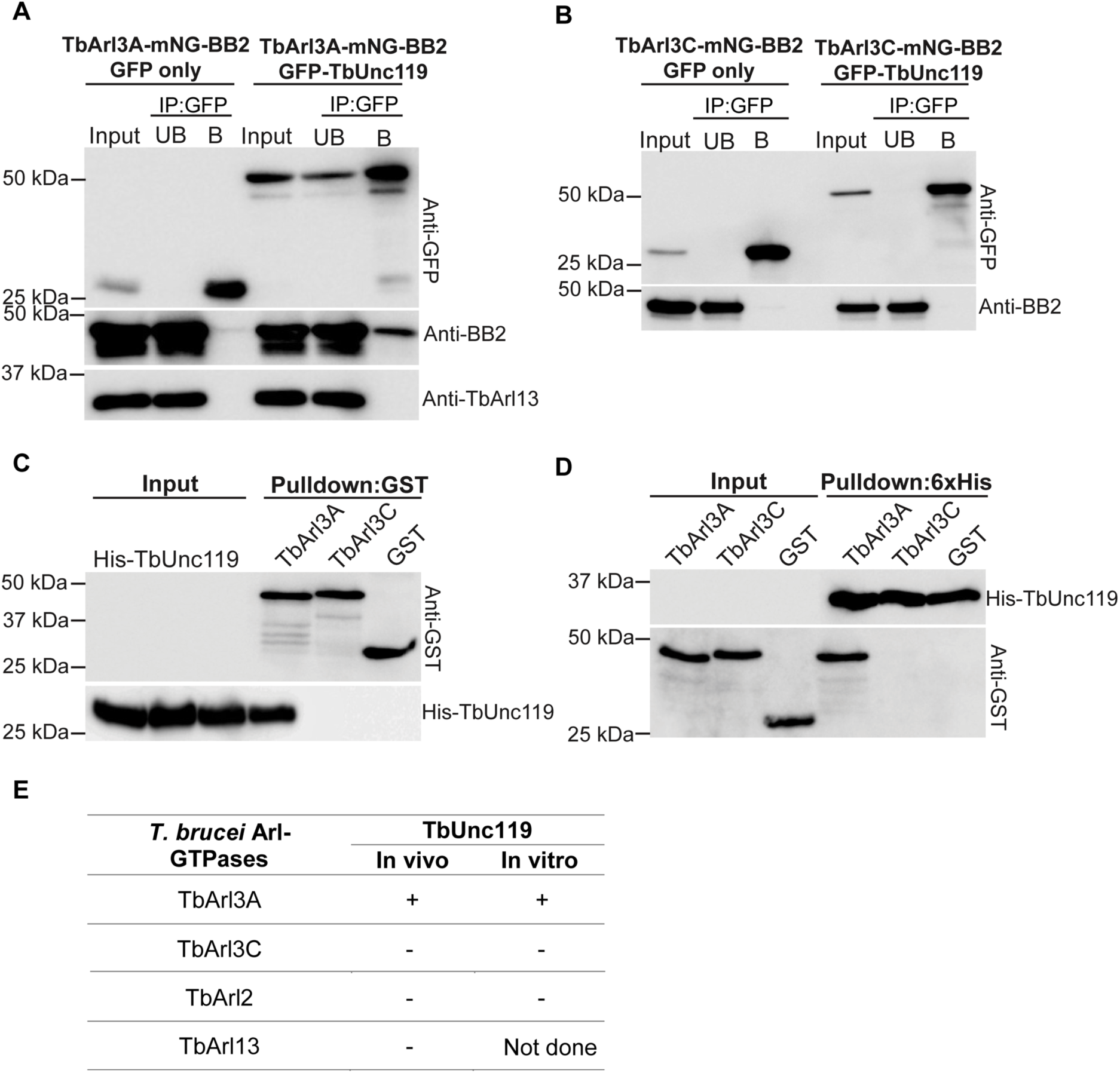
TbUnc119 interacts with TbArl3A but not TbArl3C. (**A, B**) *T. brucei* cells co-expressing GFP-TbUnc119 and TbArl3A-mNG-BB2 (A) or TbArl3C-mNG-BB2 (B) were homogenized, incubated with GFP-nAb beads and examined for co-immunoprecipitation. Cells expressing GFP only were used as negative controls. Input: 5% of cell lysates; UB: 5% of unbound fraction; B: proteins bound to GFP-nAb beads. (**C**) Glutathione beads coated with GST-TbArl3A, GST-TbArl3C or GST only were incubated with *E. coli* cell lysates expressing His-TbUnc119. **(D**) Inverse pulldown using His-TbUnc119-coated beads. Input: 12% of cell lysates. (**E**) Summary of interactions between *T. brucei* Arl-GTPases and TbUnc119.

**Fig. 6.**
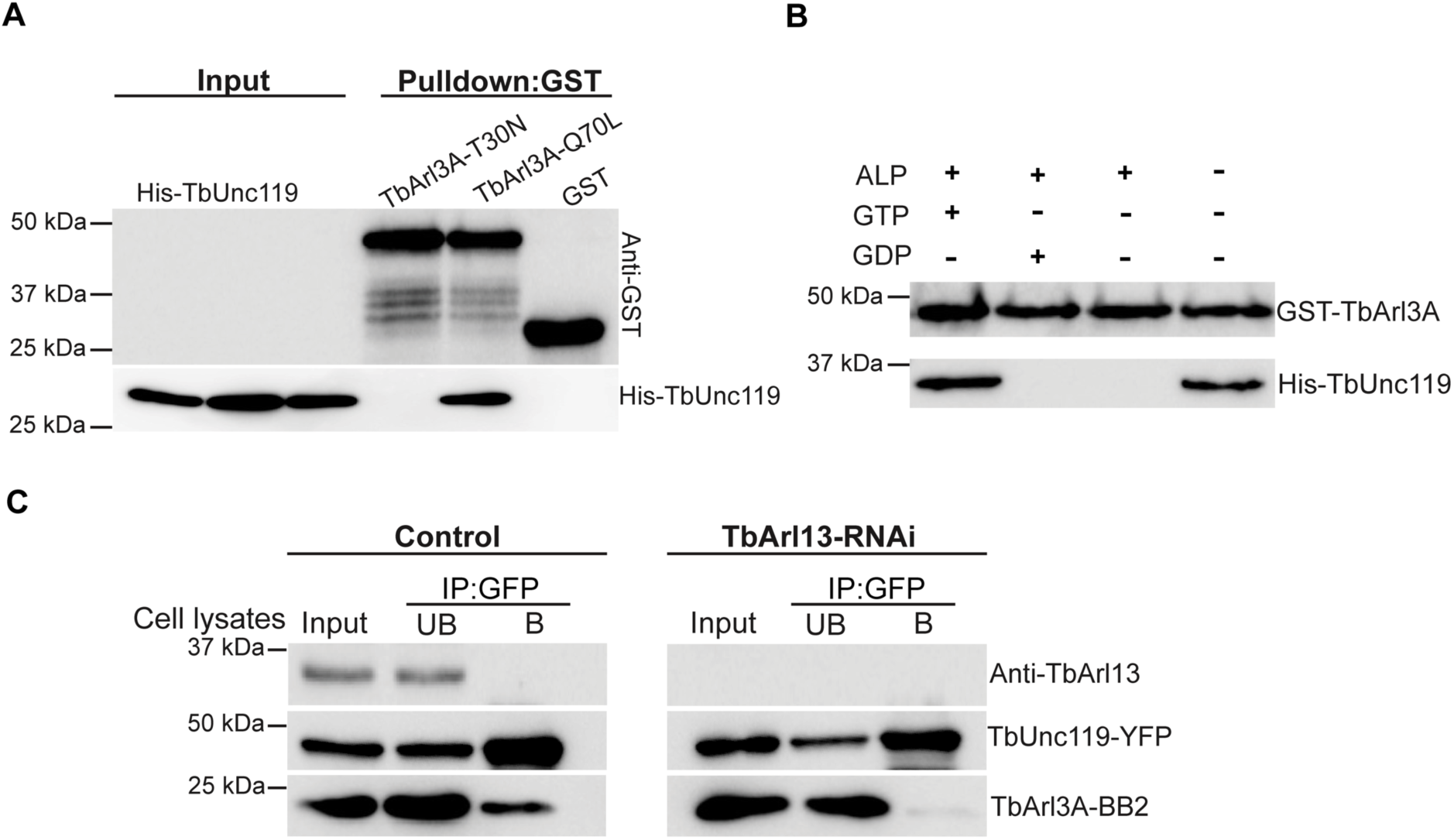
TbUnc119 interacts with TbArl3A in a GTP-dependent manner. (**A**) Glutathione beads coated with GST-TbArl3A-T30N, GST-TbArl3A-Q70L or GST were incubated with bacterial lysates expressing His-TbUnc119. Input: 12% of cell lysates. **(B**) GST-TbArl3A was treated with or without alkaline phosphatase (ALP) and then loaded with GTP or GDP was used to pull down His-TbUnc119. (**C**) Control and TbArl13-RNAi cells co-expressing TbUnc119-YFP and TbArl3A-BB2 were immunoprecipitated with GFP-nAb beads. TbArl13-RNAi did not affect TbUnc119-YFP and TbArl3A-BB2 expression levels, but their binding was abolished. Input: 5% of cell lysates; UB: 5% of unbound fraction; B: proteins eluted from GFP-nAb beads.

GTP-locked GST-TbArl3A-Q70L mutant, but not GDP-locked GST-TbArl3A-T30N could pull down His-TbUnc119 (Fig. 6A), suggesting that TbUnc119-TbArl3A interaction is GTP-dependent. This was further validated using nucleotide exchange assays (Fig. 6B). Alkaline phosphatase-treated GST-TbArl3A loaded with or without GDP did not interact with His-TbUnc119. GTP loaded TbArl3A, however, exhibited a strong and specific interaction with TbUnc119.

Our previous studies have shown that TbArl13 acts as a GEF on TbArl3A, Our previous studies have shown that TbArl13 acts as a GEF on TbArl3A,as has been reported in mammals (3,24). We hypothesized that in cells depleted of TbArl13, the level of TbArl3A-GTP in *T. brucei* should decrease, which in turn will affect the interaction between TbUnc119 and TbArl3A. To test this, stable cells expressing tetracycline-inducible TbArl13-RNAi and cumate-inducible TbUnc119-YFP and TbArl3A-BB2 were generated. While TbArl3A-BB2 co-immunoprecipitated with TbUnc119-YFP in control, their interaction was abolished in TbArl13-RNAi (Fig. 6C). Together, these results demonstrate specific interaction of TbUnc119 with only one of the TbArl3 homologs, TbArl3A, in a GTP-dependent manner, suggesting that TbUnc119 is an effector of TbArl3A that is regulated by TbArl13.

## Discussion

In this study, we revisited the functions of TbUnc119 in light of recent understanding of the LIFT pathway in ciliary biogenesis. Overall our results supported a function of TbUnc119 in flagellar targeting of myristoylated TbAK3. This is consistent with the cargo carrier function of Unc119 observed in mammals and *C. elegans* (6,19). There are, however, some differences between TbUnc119 and its higher eukaryotic counterparts. In *C. elegans*, Unc119 forms mutual interactions with both Arl3 and Arl13, facilitating GTP loading to Arl3. As the interaction between Unc119 and Arl3 is GTP-independent, Unc119 is unlikely an Arl3 effector in *C. elegans* (20). In *T. brucei*, no detectable interaction was observed between TbUnc119 and TbArl13. TbUnc119 directly interacts with TbArl3A in a GTP-dependent fashion, similar to mammalian Unc119 (5,6). Thus *C. elegans* Unc119 may represent a case of functional divergence, though it appeared more conserved with mammalian Unc119 in the phylogenetic analyses (Fig. S1A). One important difference between *T. brucei* and mammalian Unc119 is the lack of Unc119-Arl2 interaction in *T. brucei.* In mammals, Arl2 is shown to interact with Unc119 and displace low-affinity cargoes in the cytosol (5). TbArl2 is essential for cytokinesis in *T. brucei* (40) but this effect is unlikely mediated by TbUnc119.

In *T. brucei*, we showed that the binding between TbUnc119 and TbAK3 depended on the myristoylation state of the cargo. TbUnc119 is able to bind to other myristoylated proteins such as TbSMP1-1, though the function of this binding remained unclear. Depletion of TbUnc119 had no obvious effects on TbSMP1-1 distribution. Considering the lack of growth phenotypes in cultured TbUnc119-RNAi PCF and BSF cells, depletion of TbUnc119 is unlikely to cause gross perturbation in myristoylated protein distribution or functions, unlike those observed in cells with the myristoylation pathway inhibited (41-43). Depletion of TbAK3, a flagellar protein identified as TbUnc119 cargo in this study, is also shown to be dispensable for cell growth in culture (34). However, TbAK3-depletion impairs cell motility and parasite infectivity in the tsetse flies (34), suggesting that TbAK3 is crucial for flagellar function and parasite development in tsetse fly. TbUnc119 is thus expected to be important for parasite survival in hosts, which remained to be tested. Furthermore, TbUnc119 has many cytoplasmic BioID candidates without predicted myristoylation modification. During the bioinformatic analyses, we could not identify a canonical homologue of PDE6δ, a prenylated cargo carrier, in *T. brucei* and most other single-cellular organisms. Yet protein prenylation and the molecular machinery is clearly present in *T. brucei* (44-46) and several other protists (47). This raised an interesting possibility that TbUnc119 and other protist Unc119 orthologs, may be able to carry other lipidated cargoes, particularly prenylated proteins. This possibility should be examined in the future as little is currently known about prenylated targets in *T. brucei*.

A single Arl3 homolog is present in mammals and *C. elegans*, and it is known to interact with and regulate IFT components in addition to its function in displacing cargoes associated with Unc119 (48,49). *T. brucei* contains three Arl3 homologs. TbArl3A and TbArl3C are highly conserved and syntenic among kinetoplastids, and TbArl3B is diverse with unknown functions (24). While both TbArl3A and TbArl3C function in flagellar biogenesis and can be regulated by TbArl13, only TbArl3A-GTP interacted with TbUnc119 and this interaction is regulated by TbArl13. While it remains to be explored if TbArl3A may have effectors other than TbUnc119, the results support the presence of a conserved LIFT pathway in *T. brucei* that involves TbArl13, TbArl3A and TbUnc119. Our results also suggest functional diversification and specialization of Arl3-GTPases in *T. brucei.* TbArl3A is localized in the flagellum and the cytoplasm, whereas TbArl3C is restricted to the basal bodies (24). They may function at different subcellular locations on different effectors, which together contribute to the essential phenotypes observed for TbArl13.

## Experimental procedures

### Bioinformatic analyses

The amino acid sequences of Unc119 and PDE6δ from various model organisms were obtained from UniProt, Tritrypdb and NCBI protein database. Multi-sequence alignments were performed using MUSCLE (50). The output of multi-sequence alignments was formatted using Multiple Align Show of the Sequence Manipulation Suite (JavaScript application) (51). For phylogenetic analyses, the Unc119 and PDE6δ sequences were aligned using MAFFT (LINSI). ProtTest (v. 3.4.2) (52) was used for model selection. Maximum likelihood tree was generated using RAxML (v. 8.2.10) and Multiparametric bootstrapping was done using automatic bootstrapping option (autoMRE).

### Expression constructs, Cell culture and Transfection

All *T. brucei* sequences used in this study were retrieved from the Tritryp database (http://tritrypdb.org/tritrypdb/). RNAi target sequences were selected using RNAit (53). The details of plasmid constructs used in this study are summarized in Table S2.

The insect-proliferative, procyclic form (PCF) of *T. brucei* cells were cultured in Cunningham medium supplemented with 10% heat-inactivated fetal bovine serum (FBS; HyClone) at 28°C. Stable transfection conditions for PCF cells were performed according to previously published protocol (54). 29.13 *T. brucei* cells genetically modified to express T7 RNA polymerase and tetracycline repressor (55) were used to generate TbUnc119-RNAi cell line. DIpAnt, a PCF *T. brucei* cell line engineered for tetracycline-inducible and cumate-inducible expressions (32,56) was used to generate stable cell lines for cumate-inducible overexpression and/or tetracycline induced knockdown experiments. For RNAi in the bloodstream form (BSF) cells, either a single marker Lister 427 cell line (57) or a double inducible DIb427 cell line (32) was used.

### Live cell imaging, immunofluorescence and microscopy

PCF cells expressing fluorescently tagged proteins were harvested, resuspended in 1 × PBS and spread on the surface of 1% low melting point agarose gel (Bio-Rad) prepared in conditioned medium. The gel portion containing the cells was excised and placed on an imaging dish with the parasite side facing the coverslip. The partially immobilized cells trapped between the agarose gel and the cover slip could be imaged at room temperature for at least 30 minutes or through an entire cell cycle with appropriate temperature and CO2 control (58).

For immunofluorescence assays, *T. brucei* cells expressing fluorescent or small tags were washed and resuspended in 1 × PBS and attached to coverslips. Cells were fixed with 4% PFA and permeabilized with 0.25% Triton X-100 unless otherwise stated. DNA was stained with DAPI (2.5 µg/ml). Images were captured by Zeiss Axio Observer Z1 fluorescence microscope with a 63× objective (NA=1.4) and a CoolSNAP HQ2 CCD camera (Photometrics).

### Image quantification and statistical analyses

For quantifications, images acquired using fixed exposure conditions were processed using ImageJ. Fluorescence intensity of the flagellum was performed using the plot profile function, by drawing a 2 µm line (width = 5 pixels) along the distal overhang of the flagellum, where it is not attached to the cell body. Fluorescence intensity of the cytosol was quantitated over a 2 µm line (width = 5 pixels) in the posterior region of the cytoplasm away from the kinetoplast and nucleus. The membrane association of TbSMP1-1-GFP or TbSMP1-1(G2A)-GFP proteins was quantitated by plotting a 5 µm line (width = 1 pixel) transverse the posterior region of the cell body, away from the kinetoplast and nucleus. The fluorescence intensity measurements were plotted on GraphPad Prism 5. The p-values were calculated using two-tailed t-test with 95% confidence interval.

### Co-immunoprecipitation and pulldown assays

0.5-1 × 108 cells co-expressing GFP-TbUnc119 (induced with 5 µg/ml cumate for 24 hrs) and BB2 tagged Arl-GTPase (39) were harvested by centrifugation at 4500 g for 7 mins at room temperature. After 2 washes with 1× PBS, the cells were resuspended in 1× PBS supplemented with protease inhibitor cocktail (Sigma) and homogenized by sonication. The cell lysates were centrifuged at 17000 g for 15 mins at 4°C and the cleared supernatants were incubated with magnetic GFP-nAb™ beads (Allele Biotechnology) to co-precipitate GFP-fusion proteins together with their binding partners. Cells co-expressing GFP and BB2 tagged TbArl-GTPase were used as negative controls. Proteins bound to the magnetic beads were eluted by boiling in 1×Laemmli buffer and analysed by SDS-PAGE followed by immunoblotting.

GST-Arl GTPases and His-TbUnc119 fusion proteins were expressed in *E. coli* (0.1mM IPTG). Purification of these tagged proteins was performed using Ni-NTA beads (Qiagen) or Glutathione Sepharose™ 4B beads (GE healthcare) according to manufacturer’s instructions. His-TbUnc119 bound to Ni-NTA beads were incubated with cell lysates containing GST-Arl3A, GST-Arl3C and GST only. Alternatively, Glutathione Sepharose™ 4B beads bound to GST or GST-fusions including GST-Arl2 were incubated with His-TbUnc119. Interaction between TbUnc119 and Arl-GTPases was then examined by SDS-PAGE followed by immunoblotting.

### Nucleotide exchange assay

GST-TbArl3A bound to glutathione beads was treated with alkaline phosphatase (ALP,10 units/ml) to enzymatically dephosphorylate purified GST-TbArl3A (containing a mixture of GTP- or GDP-bound forms) to nascent guanosine state in 1 ml exchange buffer (20 mM HEPES pH 7.4, 1mM MgCl2, 1 mM DTT) supplemented with 50 mM EDTA for one hour at room temperature. The beads were washed thrice with 1ml exchange buffer and then incubated with GTP (100 µM), GDP (100 µM) or no nucleotide, in the presence of 100 mM MgCl2 for 1 hour at room temperature. Beads were washed once each with exchange buffer and 1 × PBS and incubated with purified His-TbUnc119 for 4 hours at 4°C. Beads were then washed thrice each with 1% Triton X-100 in 1× PBS followed by 1× PBS and bound proteins eluted by boiling in 1×Laemmli buffer.

### Proximity-dependent biotinylation (BioID) and LC-MS/MS analyses

Approximately 109 cells were induced for the expression of 3HA-BioID2-TbUnc119 and TbUnc119-BioID2-HA with 5 µg/ml of cumate for 16 hours. Wild type cells were used as negative control. 50 µM of Biotin was added to each culture 8 hours prior to harvest. Cells were washed extensively with PBS, and lysed with lysis buffer (0.4% SDS, 500 mM NaCl, 5 mM EDTA, 1 mM DTT, 50 mM Tris-HCl, pH 7.4) supplemented with protease inhibitors (Sigma). Cell lysates were centrifuged, and the clear supernatant was incubated with Streptavidin-coated Dynabeads® (Invitrogen) for 4 hrs or overnight at 4°C. The beads were washed twice with PBS containing 1% SDS, twice with PBS containing 1% Triton X-100 and then twice with 1 × PBS, for 5 mins each. The bound proteins were treated on beads with triethylammonium (500 mM, pH 8.5) and reduced with 4 µl 100 mM TCEP (Tris(2-carboxyethyl)phosphine) at 50°C for 1 hour with gentle mixing. Beads were alkylated with 5 mM MMTS (methyl methanethiosulfonate) at room temperature for 15 min. After overnight on-bead digestion with 2.5 µg Trypsin, peptide fragments were desalted and processed for LC-MS/MS analyses.

LC-MS/MS was performed on Eksigent nanoLC Ultra and ChiPLC-nanoflex (Eksigent, Dublin, CA) in Trap Elute configuration. Tandem MS analysis of digested peptide samples was performed using a TripleTOF 5600 system (AB SCIEX, Foster City, CA, USA) in Information Dependent Mode. MS spectra were acquired using the Analyst 1.6 software (AB SCIEX). The MS data fed to MASCOT Server 2.4.0 (Matrix Science) were searched against the *T. brucei* protein database (*Tbrucei*TREU927, Release-28).

Proteins identified in 3HA-BioID2-TbUnc119 and TbUnc119-BioID2-HA samples but not in the negative control were ranked based on protein content calculated from EMPAI score (59) using 1% False Discovery Rate (1% FDR). Top candidates identified in 3HA-BioID2-TbUnc119 and TbUnc119-BioID2-HA samples were scanned for the presence of myristoylation consensus sequence (MGXXXS/T) as well as their presence in the previously published myristoylation proteome of *T. brucei* (36).

### Antibodies for immunostaining and immunoblots

Antibodies used for immunostaining were anti-HA (1:500; Santa Cruz, #sc-7392) and streptavidin-Alexa Fluor 568 (1:2000). For immunoblots, the samples were boiled in 1×Laemmli buffer and separated on SDS-PAGE. Antibodies used for probing were anti-YFP (1:1000, rabbit), anti-mCherry (1:500; rabbit, Thermo Fisher, #PA5-34974), anti-TbArl13 (1:2000, rabbit) (24), anti-TbBiP (1:1000, rabbit) (60), anti-TbVDAC (1:2000, rabbit) (61), anti-His (1:5000, mouse; GE Healthcare) and anti-GST (1:5000, mouse; Santa Cruz Biotechnology). Monoclonal anti-BB2 antibodies were a kind gift from Dr. Philippe Bastin.

## Acknowledgements

We would like to thank Dr. Philippe Bastin (Pasteur Institute) for the anti-BB2 antibodies, Dr. Yiliu Zhang for the TbUnc119-YFP construct and initial discussions inspiring this work, and Dr. Amrita Srivathsan for help with the phylogenetic analyses. This study is funded by a research grant from Singapore Ministry of Education (MOE2017-T2-2-109).

## Conflict of interest

The authors declare that they have no conflict of interest with the content of this article.

## Author contributions

MP and CYH designed the experiments. MP performed and analysed the biochemistry and cell biology experiments. YH generated TbArl-GTPase associated bacterial constructs and cell lines. TK and Lin Qingsong performed LS/MS-MS and helped with data analyses. MP and CYH wrote the manuscript.

## Figure legends

**Fig. S1. *T. brucei* contains a single orthologue of Unc119.** (**A**) Maximum likelihood phylogeny of TbUnc119 with other Unc119 and PDE6δ homologues. A phylogram was generated using published Unc119 (including Unc119A and Unc119B) and PDE6δ sequences found in model organisms. Bootstrap values of >=70 are shown. Unc119 or PDE6δ homologues have not been found in yeasts and land plants, which lack cilium/flagellum. (**B**) Unc119 is highly conserved in kinetoplastids. Multi-sequence alignment of kinetoplastid homologues of Unc119, ranging from the early ancestor *P. confusum* (*Paratrypanosoma confusum*) to diverging sub-genera of C. *fasciculata, Leishmania* (*L. major and L. mexicana*), *Leptomonas* (*Leptomonas seymouri*) and *Endotrypanum* (*E. monterogeii*) and different species of *Trypanosoma*. The conserved, identical residues are highlighted in green and similar residues are in blue.

**Fig. S2. TbUnc119 is not essential for the survival of *T. brucei* in culture.** TbUnc119-RNAi was induced with tetracycline in procyclic (**A**) and bloodstream form cells (**B**). Cell density was monitored by haemocytometer counting, and the doubling number was plotted for at least 120 hrs post induction. Representative results of two independent experiments are shown.

**Fig. S3. Validation of TbUnc119-BioID constructs.** (**A, B, C**) Cells with cumate-inducible expression of 3HA-BioID2-TbUnc119 or TbUnc119-BioID2-HA were fixed and stained with anti-HA (green) and streptavidin Alexa Fluor 568 (red) to label the fusion proteins and their biotinylated products, respectively. Wild type cells were used as control. Arrowheads mark the presence of weak anti-HA and streptavidin signals associated with the flagella. Scale bar: 5 µm. (**D, E**) Cells expressing 3HA-BioID2-TbUnc119 or TbUnc119-BioID2-HA were solubilised with 0.4% SDS and 1%Triton X-100, centrifuged at 17,000*g* for 20 mins. The cleared supernatant was incubated with streptavidin beads. Total cell lysates (4%), cleared supernatant (input, 1%) and proteins eluted from the streptavidin beads were fractionated by SDS-PAGE and immuno-probed with streptavidin-HRP. Wild type cells were used as control.

**Fig. S4. TbUnc119 does not show detectable interaction with TbArl2.** (**A**) Cells co-expressing GFP-TbUnc119 and TbArl2-BB2 were incubated with GFP-nAb beads and examined for co-immunoprecipitation. Input: 3% of cell lysates. Note non-specific binding of TbArl2-BB2 to the GFP-nAb beads, even in control cells expressing GFP only. (**B**) His-TbUnc119 does not interact with GST-TbArl2 in pulldown analyses. Glutathione beads coated with GST or GST-TbArl2 were incubated with *E. coli* cell lysates expressing His-TbUnc119. Input: 12% of cell lysates.

## Notes

### Competing Interest Statement

The authors have declared no competing interest.

