## Supplemental tables and figures for "Flagellar targeting of an arginine kinase requires a conserved lipidated intraflagellar transport (LIFT) pathway in *Trypanosoma brucei*"

**Table S1. High-confidence TbUnc119-BioID candidates identified in both BioID experiments.**

| Accession | (Protein content %) | | Description | Presence in Myristoylation proteome |
|---|---|---|---|---|
| | 3HA-BioID2-TbUnc119 | TbUnc119-BioID2-HA | | |
| Tb927.9.6250, Tb927.9.6210, Tb927.9.6290 | 2.67 | 2.33 | arginine kinase | Yes (Consensus) |
| Tb927.2.4580 | 2.48 | 1.44 | TbUnc119 | - |
| Tb927.9.13920,Tb927.9.13820, Tb927.9.13785, Tb927.9.13795, Tb927.9.13860 | 2.01 | 3.29 | Kinetoplastid membrane protein 11-5 | - |
| Tb927.3.4590 | 1.47 | 0.37 | hypothetical protein conserved | Yes (Non-consensus) |
| Tb927.3.5340 | 0.82 | 0.79 | Hsc70-interacting protein (Hip), putative | - |
| Tb927.10.2750 | 0.77 | 1.72 | deoxyhypusine synthase | - |
| Tb927.9.11270 | 0.74 | 0.35 | t- complex protein 1 (eta subunit), putative | |
| Tb927.6.4960 | 0.68 | 0.38 | Zinc finger CCCH domain-containing protein 47 | - |
| Tb927.10.12110 | 0.65 | 1.79 | hypothetical protein conserved | - |
| Tb927.10.14700 | 0.63 | 0.80 | hypothetical protein conserved | - |
| Tb11.v5.0178 | 0.60 | 0.80 | glutamine synthetase, putative | - |
| Tb927.4.2740 | 0.57 | 0.93 | p25-alpha, putative | - |
| Tb927.10.11760 | 0.55 | 1.59 | pumilio/PUF RNA binding protein 6 | - |
| Tb927.4.2070 | 0.53 | 0.42 | antigenic protein, putative | - |
| Tb927.4.3570,Tb927.4.3590 | 0.50 | 1.41 | translation elongation factor 1-beta, putative | - |
| Tb927.9.13990 | 0.45 | 0.82 | RNA-binding protein, putative | - |
| Tb927.9.5730 | 0.43 | 0.39 | nucleosome assembly protein-like protein | - |
| Tb927.7.2640 | 0.42 | 0.79 | cytoskeleton associated protein, putative | - |
| Tb927.10.8190 | 0.42 | 0.38 | T-complex protein 1, theta subunit, putative | - |
| Tb927.8.1950 | 0.41 | 0.77 | hypothetical protein conserved | - |

| | | | | |
|---|---|---|---|---|
| Tb927.11.740 | 0.32 | 0.92 | eukaryotic translation initiation factor 5A | - |
| Tb927.7.2190 | 0.29 | 0.38 | translocon-associated protein (TRAP), alpha subunit, putative | - |
| Tb927.6.1800 | 0.28 | 1.28 | protein phosphatase 2C, putative | Yes (Non-consensus) |
| Tb927.11.16760 | 0.26 | 0.78 | T-complex protein 1, alpha subunit, putative | - |
| Tb11.01.2730 | 0.24 | 0.76 | hypothetical protein conserved | - |
| Tb927.7.6180 | 0.13 | 0.39 | hypothetical protein conserved | - |
| Tb11.v5.0629, Tb927.3.4500 | 0.13 | 0.37 | fumarate hydratase, putative | - |
| Tb11.v5.0181, Tb11.v5.0182, Tb927.7.5000, Tb927.7.5020 | 0.13 | 0.84 | 60S ribosomal protein L19, putative | - |
| Tb927.10.13720 | 0.13 | 0.37 | RNA-binding protein 29, putative | - |
| Tb927.2.5160 | 0.13 | 0.36 | chaperone protein DnaJ, putative | - |
| Tb927.9.6560 | 0.12 | 1.64 | NAK family pseudokinase, putative | Yes (Non-consensus) |
| Tb927.9.12070 | 0.11 | 0.72 | hypothetical protein conserved | - |

**Table S2. Plasmids used in this study**

| Expression type | Name | Selection Drugs *T. brucei*/*E.coli* | Insert information |
|---|---|---|---|
| **RNAi** | p2T7-TbUnc119 (tetracycline) | Phleomycin/Ampicillin | XbaI-TbUnc119 (68bp-536bp)-HindIII |
| **Endogenous expression** | pPOT v7-TbAK3-mNeonGreen | Neomycin/Ampicillin | TbAK3-mNeonGreen |
| **Inducible expression** | pDEX-GFP-TbUnc119 (cumate) | Phleomycin/Ampicillin | HindIII-GFP-XbaI-TbUnc119-BamHI |
| | pDEX-TbUnc119-YFP (cumate) | Blasticidin / Ampicillin | HindIII-TbUnc119-YFP-BamHI |
| | pDEX777-3HA-BioID2-TbUnc119 (cumate) | Neomycin/Ampicillin | HindIII-3HA-XbaI-BioID2-AvrII-TbUnc119-BamHI |
| | pDEX777- TbUnc119-BioID2-HA (cumate) | Blasticidin / Ampicillin | HindIII-TbUnc119-XbaI-BioID2-HA |
| | pDEX-TbAK3-BB2 (cumate) | Neomycin/Ampicillin | HindIII-TbAK3-XbaI-BB2-BamHI |
| | pDEX-TbSMP1-1-GFP (cumate) | Neomycin/Ampicillin | HindIII-TbSMP1-1-XbaI-GFP-BamHI |
| | pDEX-TbSMP1-1(G2A)-GFP (cumate) | Neomycin/Ampicillin | HindIII-TbSMP1-1(G2A)-XbaI-GFP-BamHI |
| | pDEX-TbArl2-BB2 (tetracycline) | Blasticidin /Ampicillin | HindIII-TbArl2-XbaI-AvrII-BB2-BamHI |
| **Constitutive expression** | pXS2-TbAK3-BB2 | Neomycin/Ampicillin | HindIII-TbAK3-NheI-BB2-BamHI |
| | pXS2-TbAK3(G2A)-BB2 | Neomycin/Ampicillin | HindIII-TbAK3(G2A)-NheI-BB2-BamHI |
| | pXS2-TbAK1-BB2 | Neomycin/Ampicillin | HindIII-TbAK1-NheI-BB2-BamHI |
| | pXS2-TbSMP1-1-BB2 | Neomycin/Ampicillin | HindIII-TbSMP1-1-NheI-BB2-BamHI |
| | pXS2-TbAK3-YFP | Blasticidin /Ampicillin | HindIII-TbAK3-NheI-YFP-BamHI |
| | pXS2-TbAK3(G2A)-YFP | Blasticidin /Ampicillin | HindIII-TbAK3(G2A)-NheI-YFP-BamHI |
| **Bacterial expression** | pET30a+-6XHis-TbUnc119 | --/Kanamycin | 6XHis-BamHI-TbUnc119-NotI |
| | pGEX-6P-1-TbArl3A-Q70L | --/Ampicillin | GST-EcoRI-TbArl3A-Q70L-NotI |
| | pGEX-6P-1-TbArl3A-T30N | --/Ampicillin | GST-EcoRI-TbArl3A-T30N-NotI |
| | pGEX-6P-1-TbArl2 | --/Ampicillin | GST-EcoRI-TbArl2-NotI |

**Fig. S1.** ***T. brucei*** **contains a single orthologue of Unc119.** (**A**) Maximum likelihood phylogeny of TbUnc119 with other Unc119 and PDE6δ homologues. A phylogram was generated using published Unc119 (including Unc119A and Unc119B) and PDE6δ sequences found in model organisms. Bootstrap values of >=70 are shown. Unc119 or PDE6δ homologues have not been found in yeasts and land plants, which lack cilium/flagellum. (**B**) Unc119 is highly conserved in kinetoplastids. Multi-sequence alignment of kinetoplastid homologues of Unc119, ranging from the early ancestor *P. confusum* (*Paratrypanosoma confusum*) to diverging sub-genera of C. *fasciculata, Leishmania* (*L. major and L. mexicana*), *Leptomonas* (*Leptomonas seymouri*) and *Endotrypanum* (*E. monterogeii*) and different species of *Trypanosoma*. The conserved, identical residues are highlighted in green and similar residues are in blue.

A

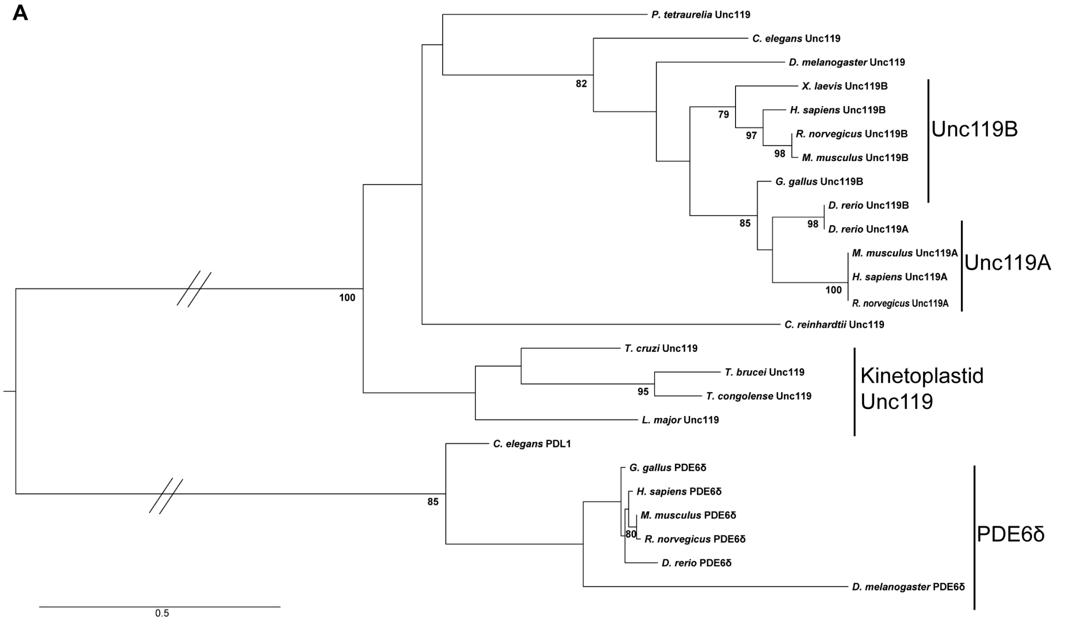


B

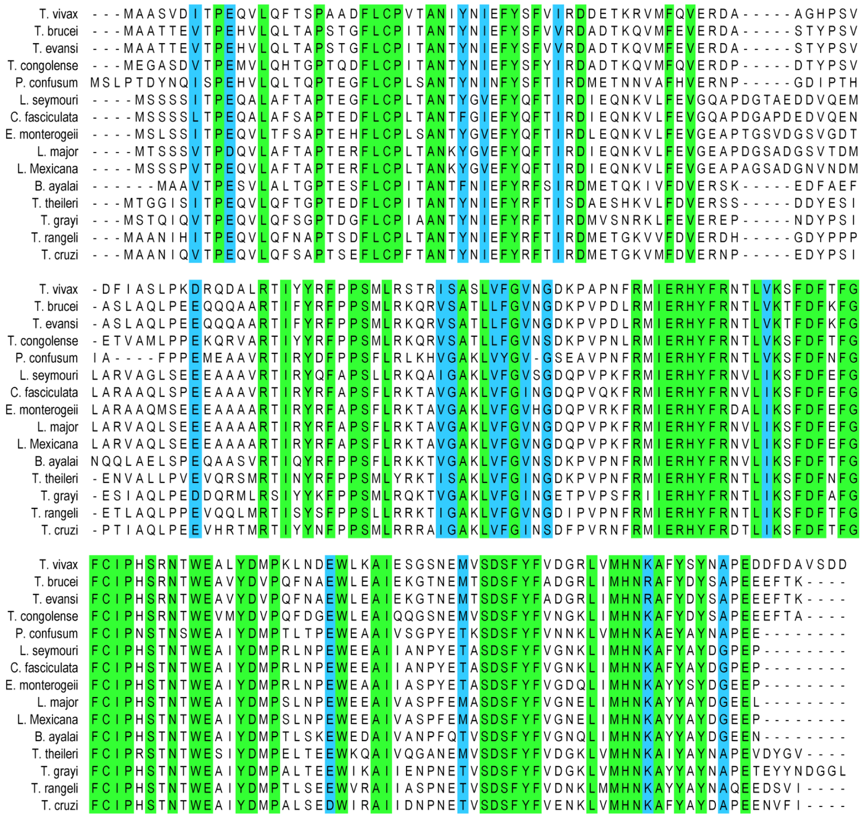

**Fig. S2. TbUnc119 is not essential for the survival of *T. brucei* in culture.** TbUnc119-RNAi was induced with tetracycline in procyclic (**A**) and bloodstream form cells (**B**). Cell density was monitored by haemocytometer counting, and the doubling number was plotted for at least 120 hrs post induction. Representative results of two independent experiments are shown.

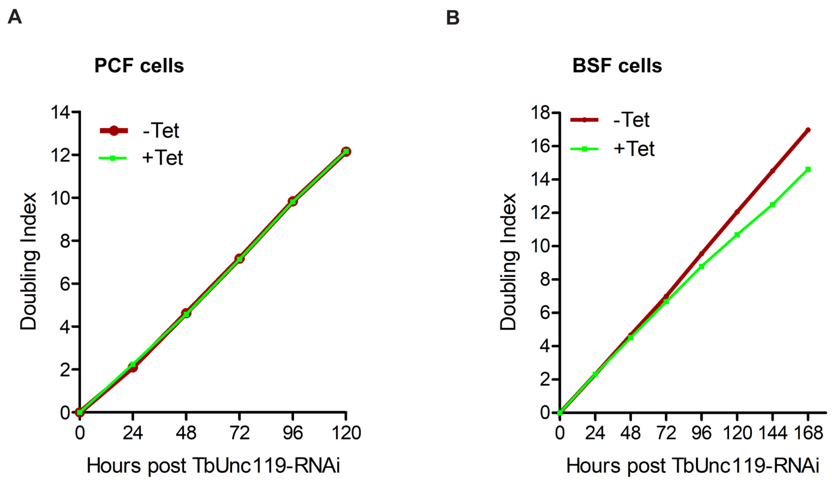

**Fig. S3. Validation of TbUnc119-BioID constructs.** (**A, B, C**) Cells with cumate-inducible expression of 3HA-BioID2-TbUnc119 or TbUnc119-BioID2-HA were fixed and stained with anti-HA (green) and streptavidin Alexa Fluor 568 (red) to label the fusion proteins and their biotinylated products, respectively. Wild type cells were used as control. Arrowheads mark the presence of weak anti-HA and streptavidin signals associated with the flagella. Scale bar: 5 µm. (**D, E**) Cells expressing 3HA-BioID2-TbUnc119 or TbUnc119-BioID2-HA were solubilised with 0.4% SDS and 1%Triton X-100, centrifuged at 17,000*g* for 20 mins. The cleared supernatant was incubated with streptavidin beads. Total cell lysates (4%), cleared supernatant (input, 1%) and proteins eluted from the streptavidin beads were fractionated by SDS-PAGE and immuno-probed with streptavidin-HRP. Wild type cells were used as control.

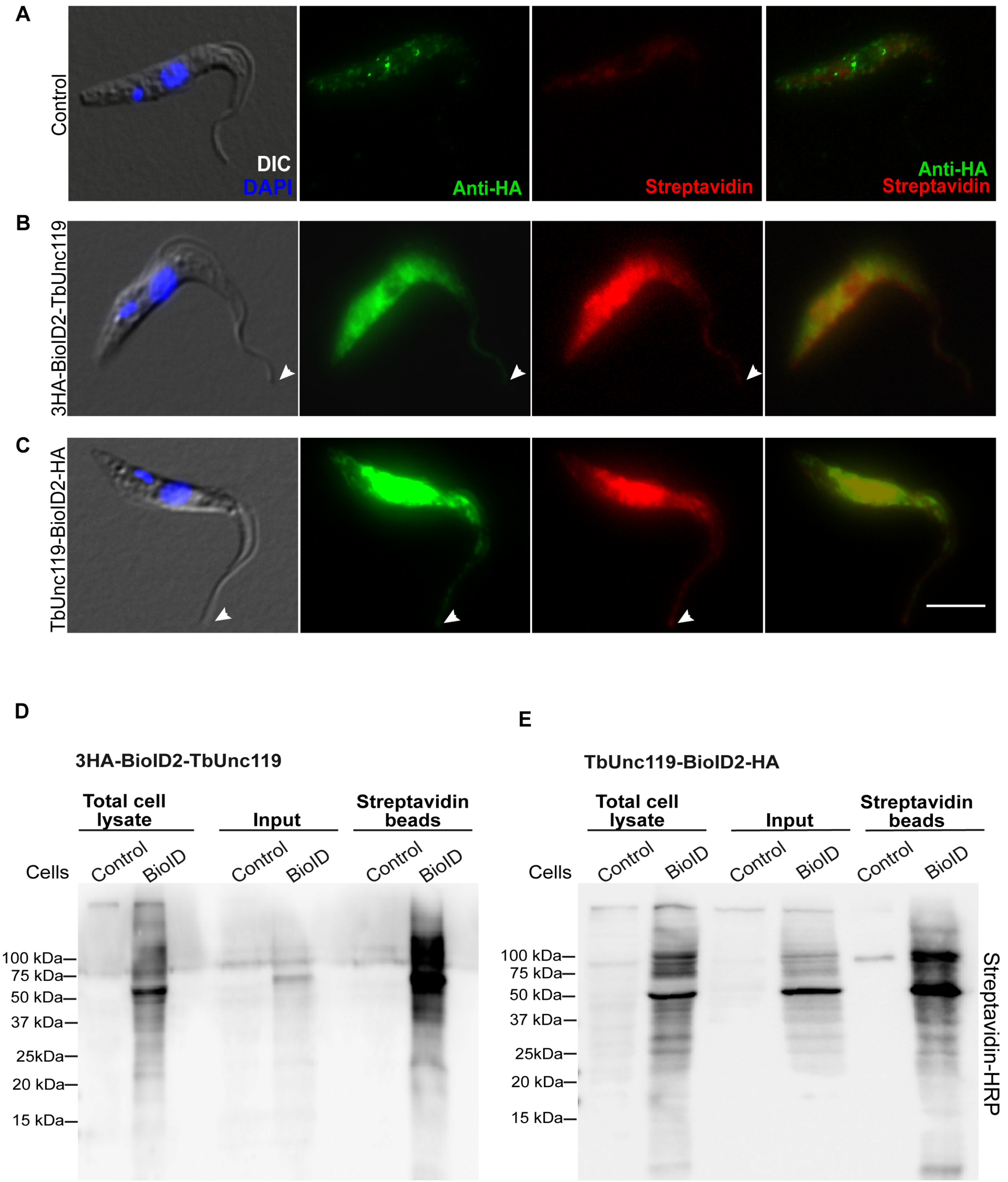

**Fig. S4. TbUnc119 does not show detectable interaction with TbArl2.** (**A**) Cells co-expressing GFP-TbUnc119 and TbArl2-BB2 were incubated with GFP-nAb beads and examined for co-immunoprecipitation. Input: 3% of cell lysates. Note non-specific binding of TbArl2-BB2 to the GFP-nAb beads, even in control cells expressing GFP only. (**B**) His-TbUnc119 does not interact with GST-TbArl2 in pulldown analyses. Glutathione beads coated with GST or GST-TbArl2 were incubated with *E. coli* cell lysates expressing His-TbUnc119. Input: 12% of cell lysates.

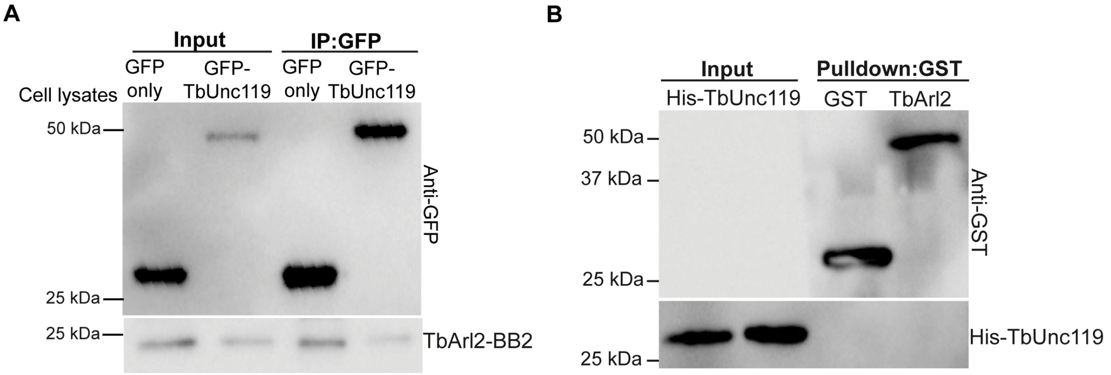
